# HLA RNAseq reveals high allele-specific variability in mRNA expression

**DOI:** 10.1101/413534

**Authors:** Tiira Johansson, Dawit A. Yohannes, Satu Koskela, Jukka Partanen, Päivi Saavalainen

## Abstract

The HLA gene complex is the most important, single genetic factor in susceptibility to most diseases with autoimmune or autoinflammatory origin and in transplantation matching. The majority of the studies have focused on the huge allelic variation in these genes; only a few studies have explored differences in expression levels of HLA alleles. To study the expression levels of HLA alleles more systematically we utilised two different RNA sequencing methods. Illumina RNAseq has a high sequencing accuracy and depth but is limited by the short read length, whereas Oxford Nanopore’s technology can sequence long templates, but has a poor accuracy. We studied allelic mRNA levels of HLA class I and II alleles from peripheral blood samples of 50 healthy individuals. The results demonstrate large differences in mRNA expression levels between HLA alleles. The method can be applied to quantitate the expression differences of HLA alleles in various tissues and to evaluate the role of this type of variation in transplantation matching and susceptibility to autoimmune diseases.

**Author Summary:** Even though HLA is widely studied less is known of its allele-specific expression. Due to the pivotal role of HLA in infection response, autoimmunity, and transplantation biology its expression surely must play a part as well. In hematopoietic stem cell transplantation the challenge often is to find a suitable HLA-matched donor due to the high allelic variation. Classical HLA typing methods do not take into account HLA allele-specific expression. However, differential allelic expression levels could be crucial in finding permissive mismatches in order to save a patient’s life. Additionally, differential HLA expression levels can lead into beneficial impact in viral clearance but also undesirable effects in autoimmune diseases. To study HLA expression we developed a novel RNAseq-based method to systematically characterize allele-specific expression levels of classical HLA genes. We tested our method in a set of 50 healthy individuals and found differential expression levels between HLA alleles as well as interindividual variability at the gene level. Since NGS is already well adopted in HLA research the next step could be to determine HLA allele-specific expression in addition to HLA allelic variation and HLA-disease association studies in various cells, tissues, and diseases.

## Introduction

The highly polymorphic human leukocyte antigens (HLA) are crucial in presentation of self, non-self and tumor antigens to T cells, and play a crucial part in autoimmunity and infection responses, as well as in organ and hematopoietic stem cell transplantation (HSCT). In the thymus and bone marrow the HLA molecules presenting self-derived peptides to maturing T-and B-cells induce the central tolerance. The classical HLA genes are divided into two classes. HLA class I genes including HLA-A, HLA-B, and HLA-C are expressed on the surface of all nucleated cells, whereas the expression of class II genes; HLA-DR, HLA-DQ, and HLA-DP is restricted to professional antigen presenting cells.[1,2] Recently a few studies reported varying expression levels of HLA alleles based on the real-time polymerase chain reaction (PCR) and the mean fluorescence intensity (MFI).[3–10] The differential expression of HLA alleles has been associated with immunologically mediated diseases, such as Crohn’s disease [11] and HIV [6,12], follicular lymphoma[7], and the outcome of HSCT through the risk of graft versus host disease (GvHD)[8,9]. In fact, incompatibilities between the donor and the recipient in HSCT have made the expression differences of HLA molecules an interesting target for finding permissive mismatches. Although currently only the qualitative HLA typing is considered in donor selection, RNAseq-based techniques can be used to determine differences in HLA expression that may influence the outcome of transplantation. The differences may also be related to the susceptibility to autoimmune diseases, tumor invasion and infections.

NGS has enabled a rapid development of several novel high-throughput HLA typing methods using different sequencing platforms.[13–22] Unlike genomic DNA based applications RNA sequencing provides a comprehensive gene expression information in addition to HLA allele calling. Precise identification of HLA alleles from NGS data is challenging due to the high polymorphism and homologous nature of HLA genes leading often to ambiguous typing results. Several existing tools, such as seq2HLA[23], HLAforest[24], and HLAProfiler[25], have been developed to perform HLA typing from short RNA sequencing reads using the whole transcriptome data. Even though these tools enable accurate and comprehensive allele determination, they only accept data with a very low error rate and are designed merely for short-read Illumina data. Owing to the complex nature of HLA genes and consequent challenges in allele assignment, ONT’s single-molecule sequencing technology has been of great interest due to its fitness for sequencing long reads.[26–28]

Here we describe a highly multiplexed RNA-based HLA sequencing method that is based on the Illumina and ONT platforms. For an accurate, high throughput quantification of the expression levels of HLA genes and alleles we developed an informatics pipeline, written in R, based on counting of unique molecular identifiers (UMI)[29,30] which work as molecular barcodes in distinguishing original transcripts from PCR copies.

## Results

We tested two different sequencing platforms, ONT and Illumina to determine HLA gene-and allele-specific expression. For this we developed a targeted ONT-based RNAseq protocol for 13 HLA genes and compared it with our Illumina-based RNAseq approach (S1 Fig). Our dataset involved RNA samples from peripheral blood of 50 healthy individuals and it consisted of 50 different HLA class I alleles and 61 different HLA class II alleles (at 2-field level) with loci HLA-B, -C and -DRB1 showing the highest heterozygosity rates of 94%, 92% and 90% respectively. The heterozygosity rate of HLA-A, -DQA1, -DQB1, -DPA1 and -DPB1 were 62%, 84%, 88%, 78%, respectively. Lower heterozygosity rates were observed with loci HLA-DPA1 (22%) and -DRA (16%). The heterozygosity rates of DRB5, and -DRB3, were 5%, and 3%, whereas all -DRB4 alleles were either homozygous or hemizygous.

### Comparison of HLA expression quantification between datasets

For accurate HLA expression analysis we determined the numbers of HLA gene- and allele-specific unique UMIs. To take into account only the unique transcripts we counted UMIs for a given gene using the UMI tools pipeline with Illumina cDNA data. To collect the number of UMIs per gene and allele, all three datasets: ONT, Illumina cDNA, and Illumina HLA amplicon, underwent the UMI counting using the custom pipeline. For the cDNA this was done to overcome the poor alignment result of HLA alleles due to the missing allelic diversity in the human reference genome. Highly homologous sequences between HLA alleles and loci made the read assignment between alleles ambiguous in some cases. The problem with multimapping reads caused by this high sequence similarity, was clear when we compared the alignment rates in the three datasets between the number of all aligning reads per HLA gene and the sum of uniquely aligning reads to the two alleles after the read assignment step. This comparison across all alleles in the Illumina cDNA showed that in average 12% (range 0.1–64%) of all reads aligning per gene were aligned uniquely to the two alleles of the gene in question. The same rates for Illumina HLA amplicon and ONT data were 48% (range 0.08–95%) and 43% (1.8–98%), respectively. The UMI duplication rate was calculated for every allele using the number of unique UMIs. Uniquely aligning reads varied in the Illumina cDNA data between 0% and 63% with the mean value of 12.6%. In the Illumina HLA amplicon data the mean duplication rate was 18.9%, (range 0% to 79%) and in the ONT data 16.5% with a range of 0–96%.

To test the correlation between the datasets, we calculated the allele-to-allele ratio from unnormalized unique UMIs for each allele pair within all 50 samples and compared the ratios to those from the Illumina cDNA and Illumina HLA amplicon data. The Illumina cDNA and Illumina amplicon data were strongly correlated (r = 0.8, p < 0.0001; Spearman rank correlation) with all HLA genes (Fig 1A), suggesting that both datasets alone were able to identify the expression difference between the two alleles. In this comparison between the two datasets, the correlation of HLA class I genes was higher (r = 0.92, p < 0.0001) compared to HLA class II genes (r = 0.69, p < 0.0001) (Fig 1B–C). In a gene-wise comparison, the strongest correlation was seen in HLA-A (r = 0.91, p < 0.0001), and HLA-B (r = 0.93, p < 0.0001) of the class I genes and HLA-DPA1 (r = 0.99, p < 0.0001), and HLA-DPB1 (r = 0.78, p < 0.0001) of the class II genes (Fig 1D–K).

**Fig 1.**
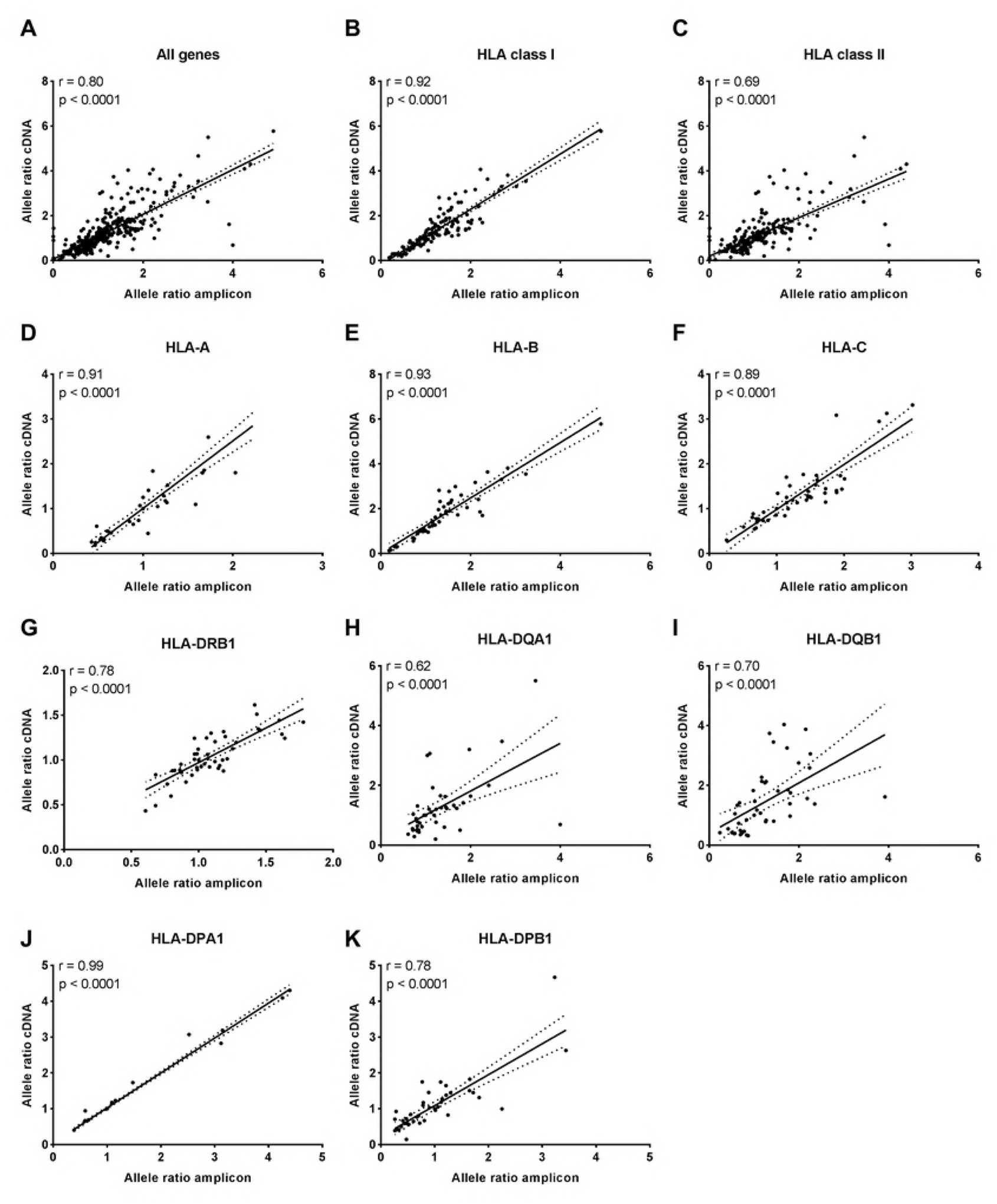
Illumina cDNA and Illumina amplicon datasets show a high correlation in allelic mRNA expression. The allele expression ratio was calculated for each allele pair in the two datasets and a non-parametric Spearman’s rank correlation was used to compare the allele-level expression between cDNA and amplicon data. Each dot represents a ratio value of heterozygous allele pairs. Homozygous allele pairs receive a ratio value of 1 which is plotted twice, once for each dataset. The line indicates the linear regression and dashed lines the 95 % confidence intervals. The Spearman correlation coefficient is given for all genes (A), HLA class I (B), HLA class II (C), and for genes HLA-A (D), HLA-B (E), HLA-C (F), HLA-DRB1 (G), HLA-DQA1 (H), HLA-DQB1 (I), HLA-DPA1 (J), and HLA-DPB1 (K). The comparison between loci DRA, DRB3, DRB4, and DRB5 is not shown due to a low number of data points.

To test the correlation between ONT and Illumina HLA amplicon data at allele level we calculated the allele ratio from ONT data as well. This comparison showed a weaker correlation with all HLA genes included (r = 0.47, p < 0.0001) (Fig 2A). The class I genes showed a moderate to strong correlation (r = 0.67, p < 0.0001), whereas the correlation of class II genes was weaker (r = 0.32, p < 0.0001) (Fig 2B–C). HLA-B (r = 0.61, p < 0.0001) and HLA-C (r = 0.79) correlated better than HLA-A (r= 0.49, p = 0.0008) (Fig 2D–F). In class II genes HLA-DRB1 (r = 0.62, p < 0.0001) and HLA-DPA1 (r =0.53, p = 0.0003) showed the strongest correlation, while the other class II genes showed a weak correlation (Fig 2G–K). Surprisingly, the same comparison between ONT and Illumina cDNA data correlated better with all HLA genes (r = 0.53, p < 0.0001) (Fig3A). Similarly, HLA class I gave a stronger correlation (r = 0.59, p < 0.0001) when compared to class II (r = 0.48, p < 0.0001) (Fig 3B–C), however, for both the correlation was moderate at best. In the gene-wise comparison the strongest correlations were seen in HLA-A (r = 0.57, p < 0.0001) (Fig 3D), HLA-B (r =0.59, p < 0.0001) (Fig 3E), HLA-C (r = 0.68, p < 0.0001) (Fig 3F), HLA-DQA1 (r=0.59, p < 0.0001) (Fig 3H), HLA-DQB1 (r =0.49, p = 0.0003) (Fig 3I), and HLA-DPA1 (r = 0.54, p = 0.0002) (Fig 3J), and the lowest in HLA-DRB1 (r = 0.46, p = 0.0022) (Fig 3G), and HLA-DPB1 (r = 0.47, p = 0.0009) (Fig 3K). The correlation comparisons of allele ratios between ONT and Illumina datasets suggest that we are either unable to assign all the reads properly to the correct alleles or that we miss UMIs in the UMI quantification step with ONT data, or both. This result indicates the difficulty of finding the UMI position in ONT reads compared to Illumina reads where the 10 bp UMI is always sequenced first in the beginning of read 1. Due to a moderate correlation result between ONT and Illumina, no gene-and allele-level expression comparison is shown.

**Fig 2.**
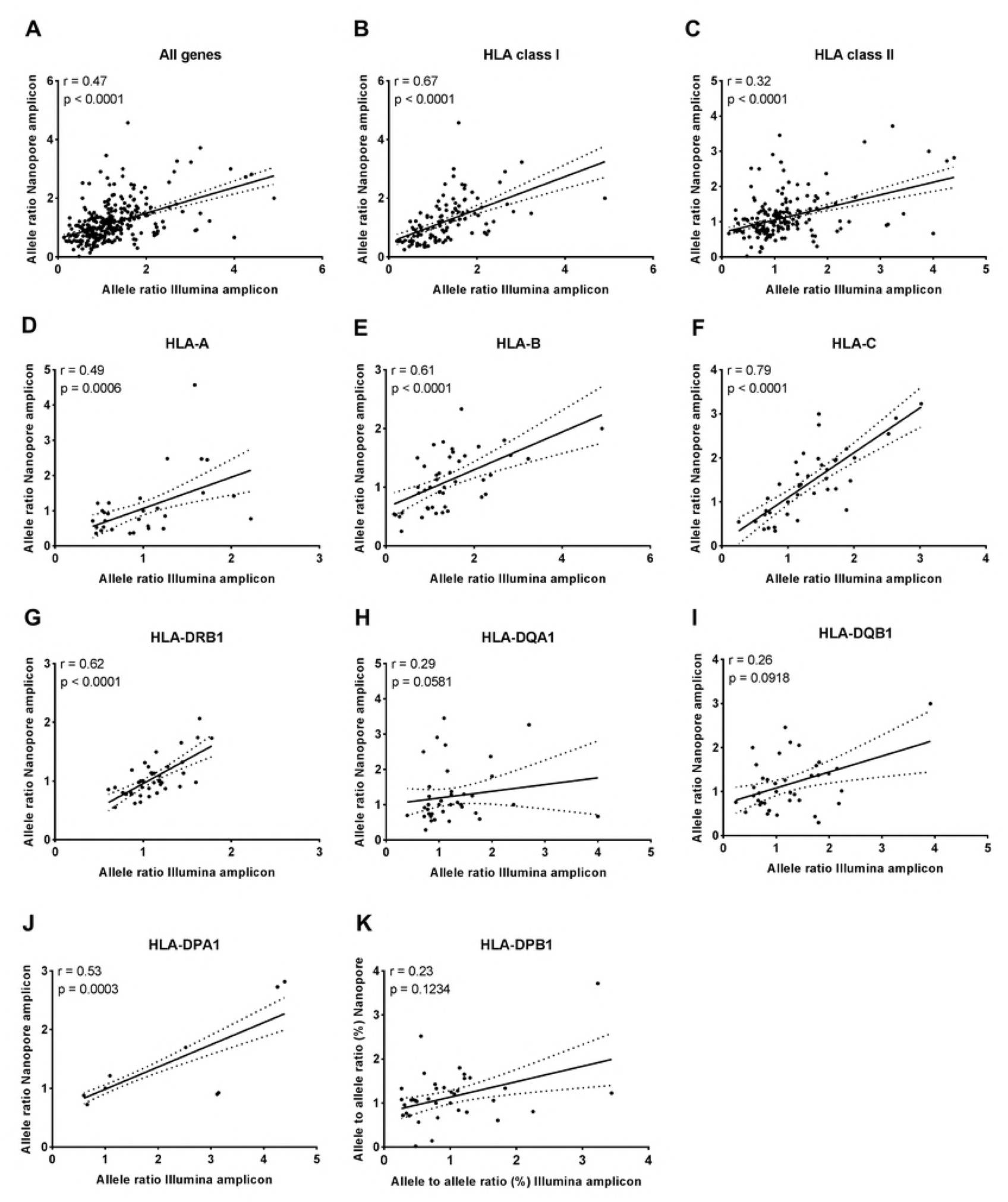
A Spearman’s rank correlation of the allele expression ratio between ONT and amplicon data shows weak to strong correlation. Correlations of allelic mRNA expression are given as expression ratios for each heterozygous allele pair which each dot represents in the scatter plot. Homozygous allele pairs receive a ratio value of 1 which is plotted twice, once for each dataset. The line indicates the linear regression and dashed lines the 95 % confidence intervals. The Spearman correlation coefficient is shown for all genes (A), HLA class I (B), HLA class II (C), and for genes HLA-A (D), HLA-B (E), HLA-C (F), HLA-DRB1 (G), HLA-DQA1 (H), HLA-DQB1 (I), HLA-DPA1 (J), and HLA-DPB1 (K).

**Fig 3.**
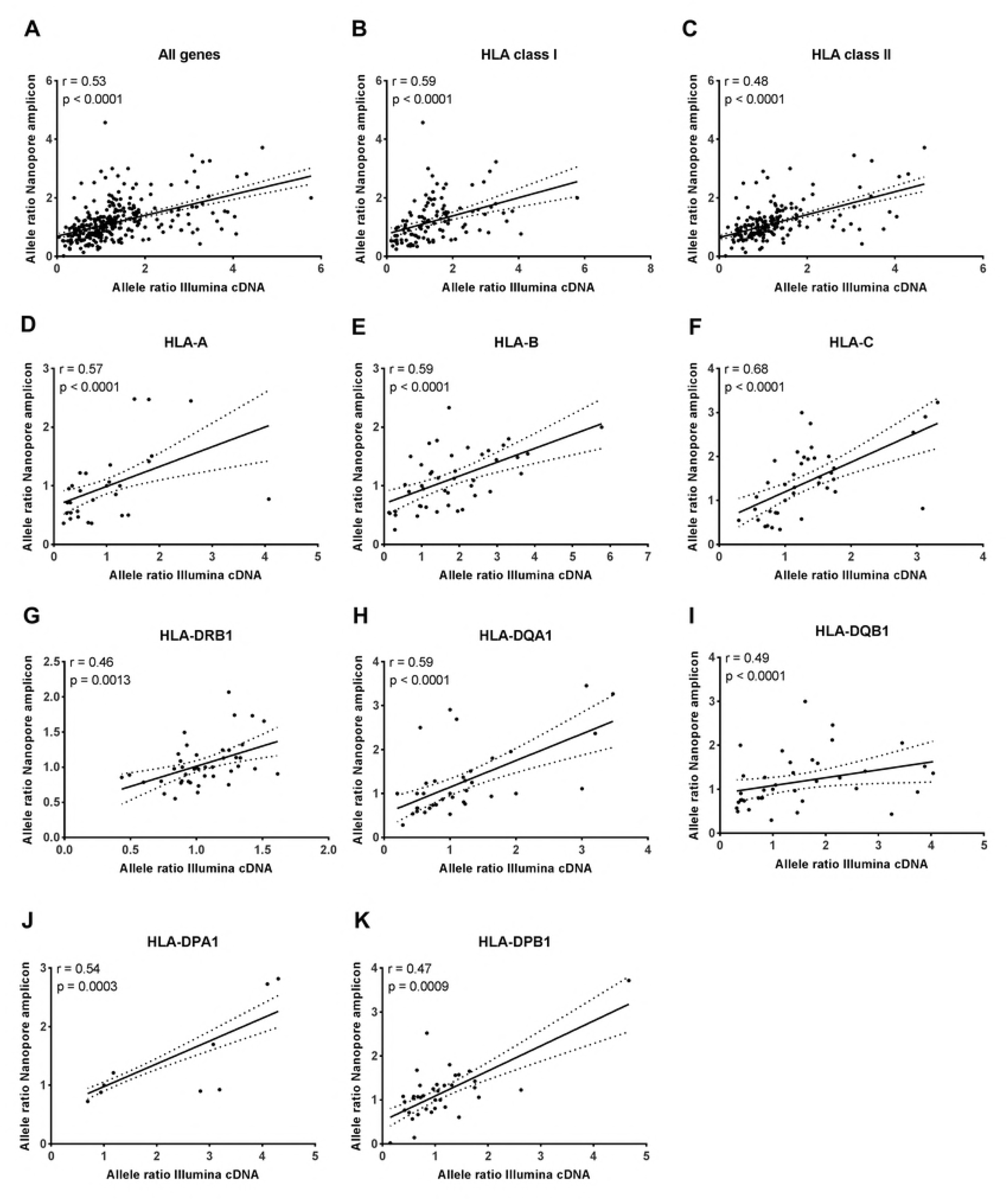
Correlation comparison of allelic HLA mRNA expression between Illumina cDNA and ONT amplicon datasets. Scatter plots showing the Spearman’s rank correlation and a linear regression of allele expression ratio between ONT and Illumina cDNA data. Dots represent a ratio value of heterozygous allele pairs. Homozygous allele pairs receive a ratio value of 1 which is plotted twice, once for each dataset. The dashed lines indicate the 95 % confidence intervals. The Spearman correlation coefficient is shown for all genes (A), HLA class I (B), HLA class II (C), and for genes HLA-A (D), HLA-B (E), HLA-C (F), HLA-DRB1 (G), HLA-DQA1 (H), HLA-DQB1 (I), HLA-DPA1 (J), and HLA-DPB1 (K).

### HLA gene-specific expression

To characterize gene and allelic expression profiles across samples Illumina cDNA and HLA amplicon UMI counts were normalized to library size using the CPM method. First, we explored the amount of HLA expression from the total expression of all genes across the samples using unique UMIs of the Illumina cDNA data. The proportion of total HLA expression out of all cDNAs varied between 0.96% and 2.54%, and HLA class I and HLA class II from 0.48% to 1.99% and 0.26% to 1.14%, respectively (S4 Fig). For the gene-level comparison the sum of two alleles was calculated from the CPM-normalized unique UMI values. This comparison was done between the Illumina cDNA and HLA amplicon datasets across the 50 samples. In Illumina cDNA data we clearly see a higher expression of HLA class I genes compared to class II, whereas in the Illumina HLA amplicon data HLA-DRB gene shows high expression values across samples (Fig 4). In the cDNA data HLA-B and -C were expressed at the highest levels. HLA-A gene expression was lower compared to the two other class I genes. In the HLA class II HLA-DRA and -DRB genes were expressed at the highest levels following -DPA1 and -DPB1. HLA-DQA1 and -DQB1 were expressed clearly at the lowest levels. The evaluation between the two Illumina datasets revealed that in the HLA amplicon dataset HLA class II has higher gene-level expression than in the Illumina cDNA dataset. The genes expressed at the highest levels in this data were HLA-DRB, and HLA class I genes. The bias towards HLA class II and especially in HLA-DRB in the HLA amplicon data most likely arises from the different efficacy rates of HLA primers used in the amplification and leading to uneven pooling in the library preparation step. Since every cell expresses HLA class I, it is logical that the expression of HLA class I genes should be higher compared to HLA-DRB expression. For this reason, in the following analyses we show the data from the Illumina cDNA dataset.

**Fig 4.**
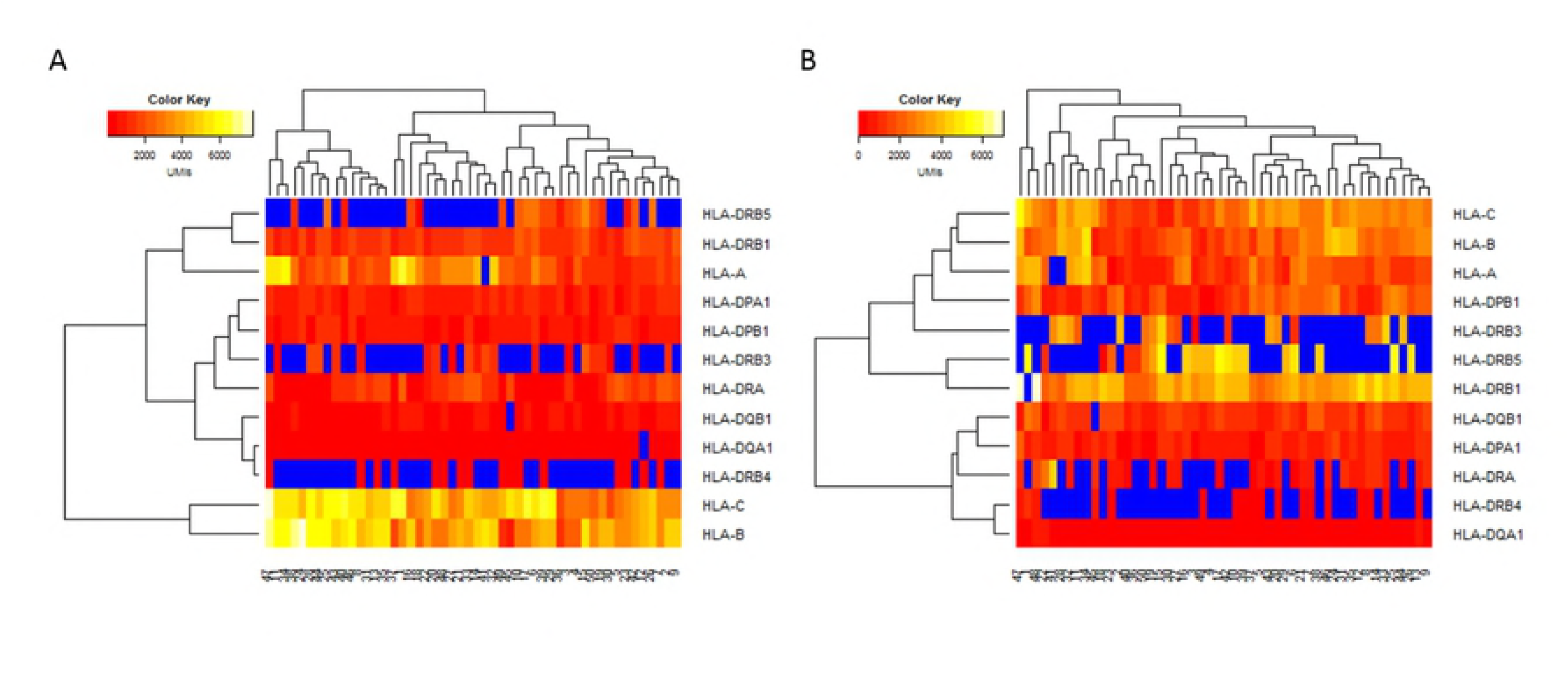
Hierarchial clustering and heatmap of gene expression levels of 12 HLA loci in the Illumina cDNA and HLA amplicon datasets. (A) The gene-specific comparison of Illumina cDNA data and (B) Illumina HLA amplicon data. The represented gene expression is the sum of unique UMIs from the two alleles (homozygous and heterozygous individuals) or the unique UMI count of on allele (hemizygous individuals) in HLA-DRB3, -DRB4, and -DRB5. The columns represent 50 individuals and the rows different HLA genes. Expression levels are colored with yellow for high expression and red for low expression. The blue color indicates missing expression values for a given gene.

The further comparison between the two Illumina datasets at the allele-specific level is shown in the supplementary information (S5–S6 Fig). The overall class-level comparison across all 50 samples showed that mRNA for HLA class I was expressed in significantly higher levels than HLA class II (p < 0.0001) (Fig 5B). Between HLA class I genes, the expression of HLA-A was lower than HLA-B (p < 0.005) and HLA-C (p < 0.005), however, there was no significant difference between HLA-B and -C mRNA expressions (Fig 5B). In the class II gene-level comparison, HLA-DR (including mRNAs for DRA, DRB1, DRB3-5) was expressed at higher level compared to HLA-DP (p < 0.0001) and HLA-DQ (p < 0.0001) (Fig 5B). The expression of HLA-DP and -DQ also differed statistically significantly (p < 0.05), the expression of HLA-DQ being the lowest.

**Fig 5.**
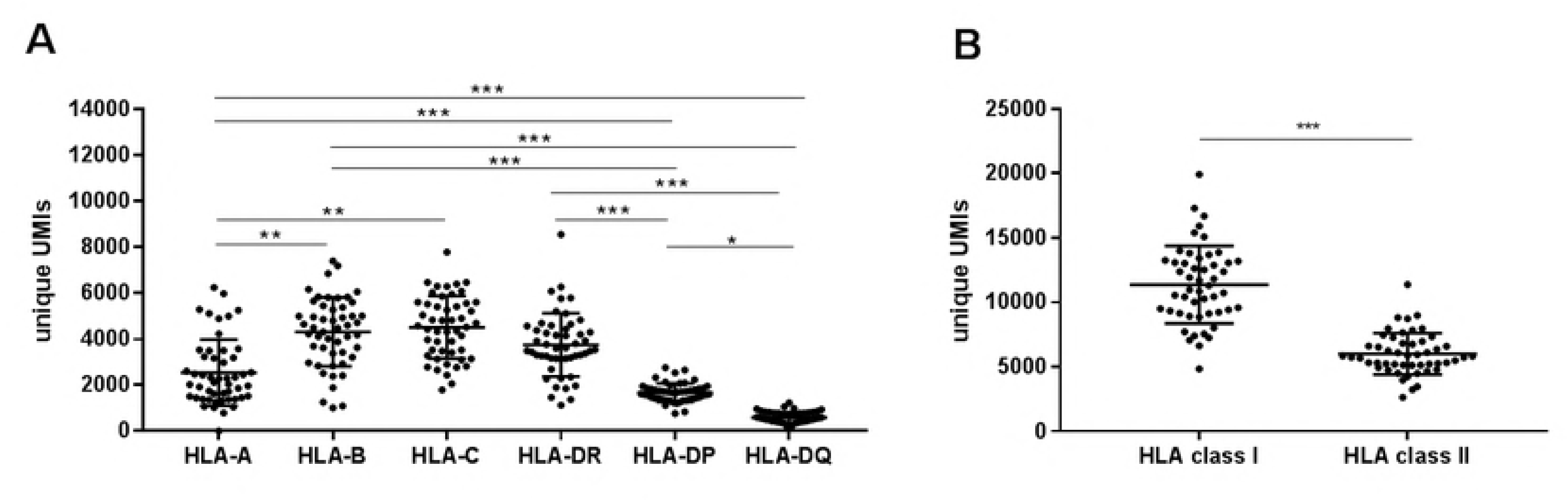
The expression of HLA class I and class II genes. (A) The mRNA expression at a heterodimer level was calculated from the allele-level unique UMIs for all 50 individuals. For class I genes the gene-specific expression corresponds to the sum of two alleles for a given gene. For HLA-DPA1/B1 and HLA-DQA1/B1 the expression value was calculated using the sum of unique UMIs from both α-and β-chain alleles (4 alleles). The expression of HLA-DR depends from the individual’s haplotype and was either calculated from the allele-level unique UMIs of HLA-DRA and HLA-DRB1 (4 alleles), or from the combination of these two and genes DRB3, DRB4, and DRB5. (B) For class-level expression comparison allele-level unique UMIs were calculated together class-wise for each individual. Each dot represents the expression value of one individual per group. Wide horizontal lines correspond to the mean expression and short horizontal lines for standard deviation for each group. A Kruskal-Wallis test was performed to compare the expression difference between HLA-A, -B, -C, -DR,-DP, and -DQ and Mann-Whitney U test to compare the expression between HLA class I and class II. *p-value < 0.05; **p-value < 0.005; ***p-value < 0.0001.

To assess the differential expression of HLA genes between individuals we calculated the relative expression of all genes present per sample using unique UMIs and compared these relative expression profiles between 50 individuals. The comparison demonstrated that the relative amounts of different HLA mRNAs varied greatly between individuals (Fig 6). In addition, the total amount of mRNA for HLA varied between individuals (data not shown). We found that in average 65% (range 45-84%) of the total HLA expression came from the HLA class I genes, whereas the average of HLA class II expression across individuals was 35% (range 16-54%).

**Fig 6.**
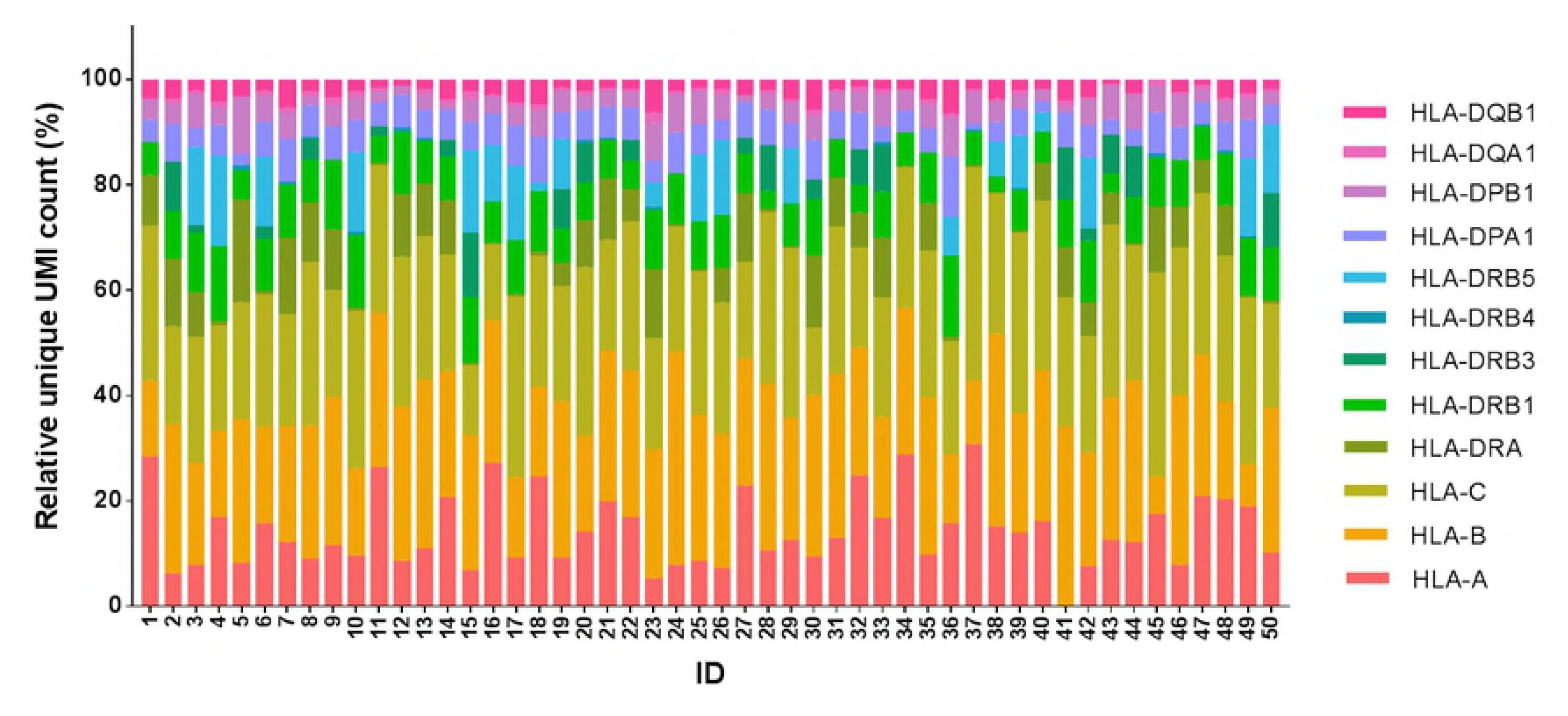
The mRNA expression distribution of 12 HLA genes across 50 individuals. The relative expression of each HLA gene was calculated from the number of unique UMIs (Illumina’s cDNA dataset) of two alleles (homozygous and heterozygous samples) or one allele (hemizygous samples) out of the total unique UMI number per individual. Different colors show the distribution of 12 HLA genes within individuals.

A comparison of HLA class I and II expression between genders (n = 27 females and n = 23 males) showed no significant difference. Also no significant correlation between the expression levels of HLA class II and the class II transactivator, CIITA (r = 0.16, p = 0.2654) was found across the 50 individuals.

### HLA allele-specific expression

To assess HLA allelic expression we studied the number of unique UMIs representing the mRNA expression of individual alleles for a given gene across all 50 samples. The mean HLA-A mRNA expression level as defined by UMIs was 1275. Compared to this level, the HLA-A alleles A*03:01 (n = 28), and A*68:01 (n = 3) had higher than the average expression levels. Alleles A*01:01 (n = 8), A*02:01 (n = 26), and A*24:02 (n = 16) were associated expression levels lower than average (Fig 7A). Alleles A*32:01 (n = 4) with a mean of 1324 was not associated to either due to their expression levels so close to the mean expression value (henceforth neutral). Homozygous allele pairs showed lower expression levels than heterozygotes in all allele groups carrying both individuals. The expression levels between different allele groups differed significantly (H = 11.75, p = 0.04), however, a pairwise comparison showed no significant differences between allele groups.

**Fig 7.**
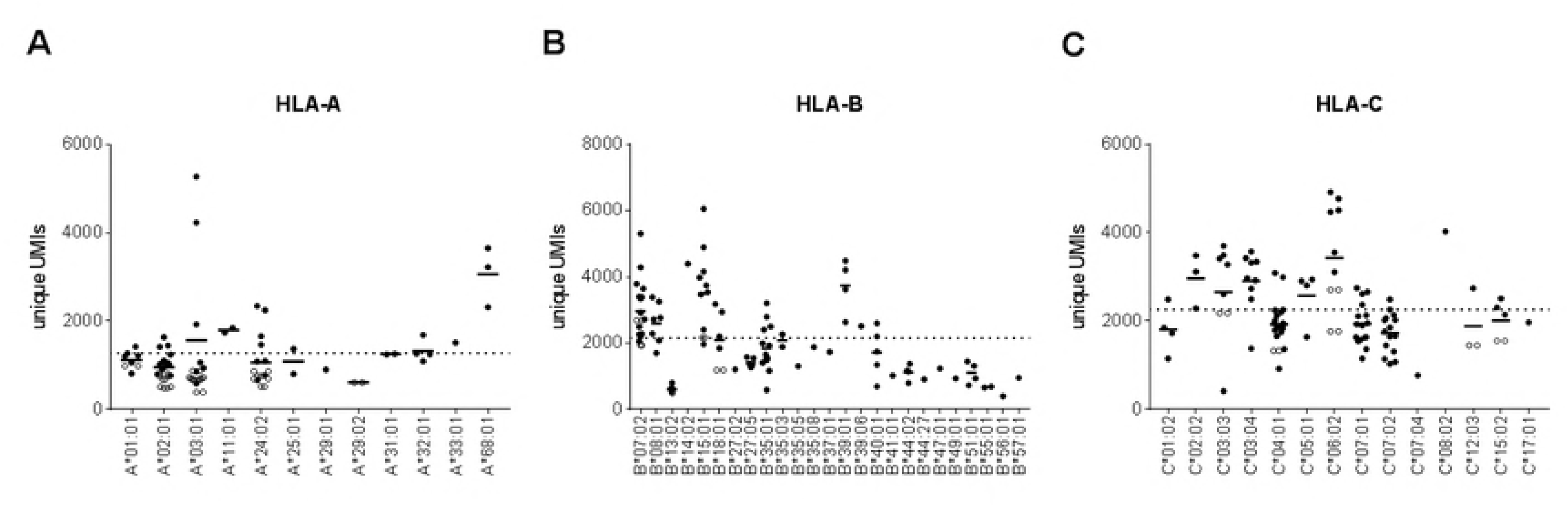
Allele-specific expression of HLA class I genes. Allele-level unique UMIs representing the allelic mRNA expression values of 50 individuals were first normalized and then grouped and plotted according to different alleles in Illumina cDNA data. Mean expression of individual alleles is indicated by a solid bar and mean expression of all alleles is represented by the dotted line. Open circles correspond to homozygous individuals. All class I genes; (A) HLA-A alleles (n = 12), (B) HLA-B alleles (n = 25), (C) HLA-C alleles (n = 14) show differential mRNA expression levels between and within allele group.

By comparing the expression levels to the mean HLA-B mRNA expression value of 2158, alleles B*07:02 (n = 18), B*08:01 (n = 7), B*15:01 (n = 11), and B*39:01 (n = 4) had a higher expression and B*13:02 (n = 6), B*27:05 (n = 5), B*35:01 (n = 14), B*40:01 (n = 5), B*44:02 (n = 4), and B*51:01 (n = 4) had a lower than the mean expression level (Fig 7B). Alleles B*18:01 (n = 6) with a mean of 2094) was considered neutral. A comparison of expression levels showed a significant difference between allele groups (H = 55.26, p < 0.0001). In the pairwise comparison significant difference (p < 0.05) was seen between pairs B*15:01∼B*44:02, B*15:01∼B*51:01, and B*39:01∼B*44:02.

Among 14 HLA-C alleles with a mean expression of 2257, C*02:02 (n = 3), C*03:03 (n = 8), C*03:04 (n = 9), C*05:01 (n = 4), and C*06:02 (n = 10) were associated with a higher expression and C*01:02 (n = 4), C*04:01 (n = 20), C*07:01 (n = 16), C*07:02 (n = 15), C*12:03 (n = 3), and C*15:02 (n= 5) with a lower expression (Fig 7C). These results correlate with previously reported allelic mRNA expression levels [3]. Similarly to HLA-A locus, we observed lower expression levels in homozygous individuals. Allele-specific expression comparison showed a significant difference between allele groups (H = 35.73, p < 0.0001). In the pairwise comparison allele groups C*03:04 ∼ C*07:02, C*04:01∼ C*06:02, and C*06:02 ∼ C*07:02 were significantly different (p < 0.05).

The comparison of HLA-DRB1 expression values to the mean expression value of 745 categorized DRB1*01:01 (n = 16), DRB1*10:01 (n = 3), and 15:01 (n = 17) into a group of high-expression associated alleles, whereas DRB1*03:01 (n = 7), DRB1*07:01 (n = 9), DRB1*13:02 (n = 5), and DRB1*16:01 (n = 4) were grouped to a low-expression (Fig 8B). Alleles DRB1*04:01 (n = 6), DRB1*08:01 (n = 10), and DRB1*13:01 (n = 12), were considered neutral. Overall, this locus was very heterozygous as only four homozygous individuals were observed in DRB1*01:01 and DRB1*08:01. In contrast to HLA-A and HLA-C, homozygous individuals in HLA-DRB1 were expressed at higher levels. The expression levels between allele groups were significantly different (H = 19.26, p = 0.02), though, no significant differences were seen between alleles in the pairwise comparison. HLA-DRA is not shown due to possible bias between homozygous and heterozygous individuals. This bias most likely results from an allele assignment problem in short Illumina reads caused by the low number of variant positions between DRA alleles. In case of a heterozygous individual carrying DRA*01:01 we constantly observed a low number of unique UMIs resulting from the second allele.

**Fig 8.**
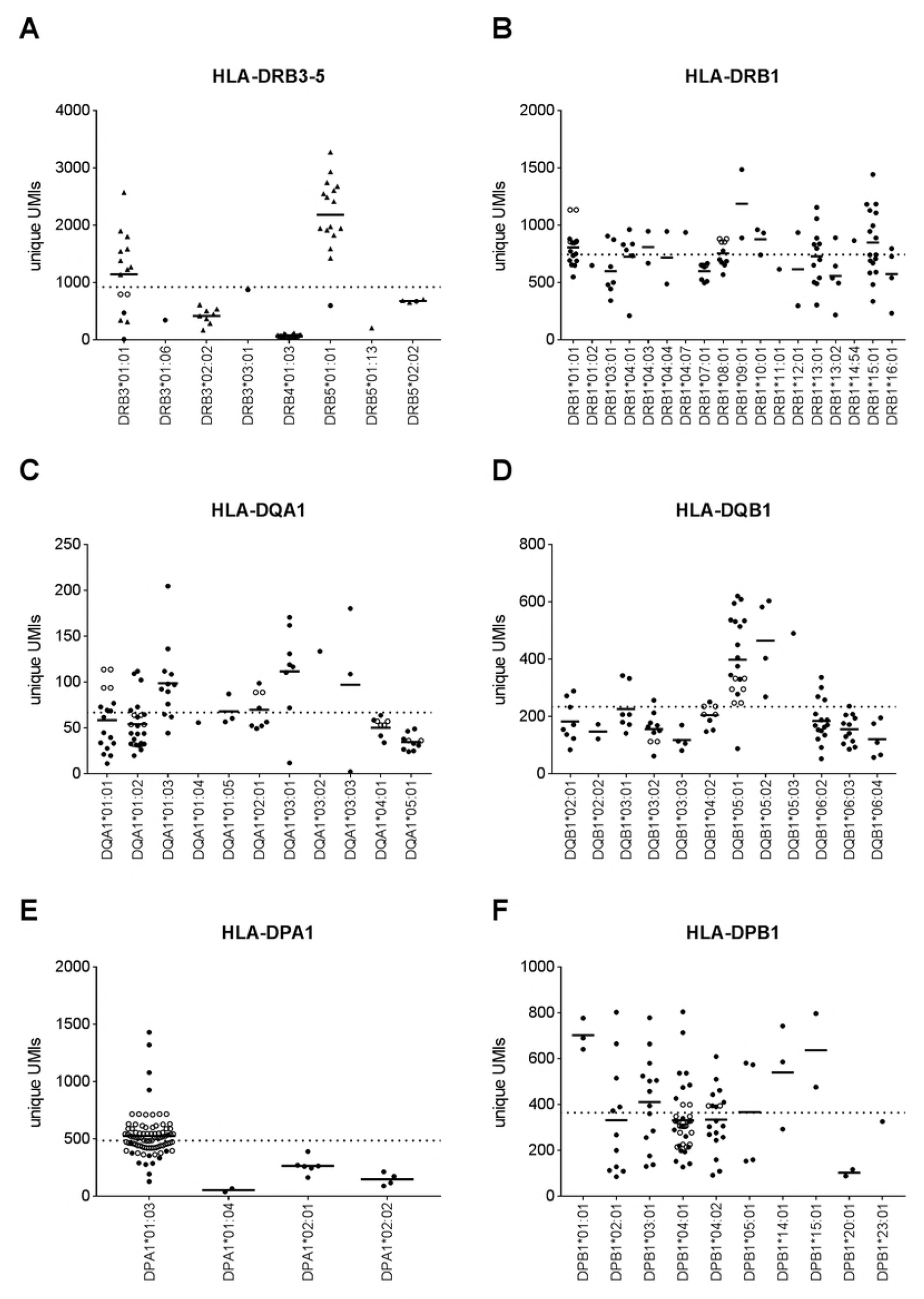
Allele-specific expression of HLA class II genes. Differential allele-specific expression profiles of 50 individuals are represented for each gene (A) HLA-DRB3 (n = 4), HLA-DRB4 (n = 1), HLA-DRB5 (n = 3), (B) HLA-DRB1 (n = 18), (C) HLA-DQA1 (n = 11), (D) HLA-DQB1 (n = 12), (E) HLA-DPA1 (n = 4), (F) HLA-DPB1 (n = 10). Each dot refers to a unique UMI value which are plotted according to alleles. The horizontal black bars indicate the mean expression of individual alleles and the dotted line corresponds to mean expression of all alleles. Open circles correspond to homozygous individuals and black triangles to hemizygous individuals (DRB3, DRB4, and DRB5).

Out of the four HLA-DRB3 alleles present in this data, DRB3*01:01 (n = 15) and DRB3*02:02 (n = 8) were the most frequent. DRB4*01:03 (n = 20) was the only allele representing this locus in our data. Among HLA-DRB5 alleles, DRB5*01:01 (n = 16) was the most frequent. In a pairwise comparison no significant differences were found between alleles. However, DRB4*01:03 was expressed at significantly lower levels than DRB3*01:01 and DRB5*01:01 (p < 0.005 for both). The majority of samples were hemizygous for DRB3, DRB4, and DRB5 and hence it was surprising that compared to the homozygotes and heterozygotes of all DRB3, DRB4, DRB5, hemizygotes were expressed at higher levels (p < 0.05) (Fig 8A). This might derive from a bias problem between two alleles in the read assignment. Reads which passed the set parameters in the read assignment after alignment are considered in the UMI counting. With homozygous and hemizygous alleles there is no need to assign reads between two alleles and hence a bias might occur if more reads are saved for the UMI counting compared to the heterozygotes.

At HLA-DQA1 locus, DQA1*01:03 (n = 12), DQA1*03:01 (n = 8), and DQA1*03:03 (n = 3) were associated with a higher expression levels when compared to the mean expression value of 67 (Fig 8C). In contrast, alleles DQA1*01:01 (n = 17), DQA1*01:02 (n = 26), DQA1*04:01 (n = 9), and DQA1*05:01 (n = 10) were linked to a lower expression. The alleles expressed at higher levels exhibited a heterogeneous expression, whereas the expression of low expression associated alleles was more uniform. Two alleles, DQA1*01:05 (n = 3) and DQA1*02:01 (n = 8) were not clearly associated to either of the former groups and hence were considered neutral. Significantly different expression levels were found between two high-low expression associated allele groups, DQA1*01:03 ∼ DQA1*05:01 and DQA1*03:01 ∼ DQA1*05:01 (p < 0.05 for both). Among HLA-DQB1 alleles, only two alleles, DQB1*05:01 (n = 20), DQB1*05:02 (n = 4) were associated with a higher expression compared to the mean expression value of 234 (Fig 8D). The other DQB1 alleles, DQB1*02:01 (n = 8), DQB1*03:02 (n= 10), DQB1*03:03 (n = 4), DQB1*04:02 (n = 9), DQB1*06:02 (n = 16), DQB1*06:03 (n = 12), and DQB1*06:04 (n = 5) were associated to a lower expression with more homogenous distribution. Allele-level expression was different between the allele groups (H = 49.21, p < 0.0001) and the pairwise comparison showed a significant difference (p < 0.05) between allele groups DQB1*03:02 ∼ DQB1*05:01, DQB1*03:03 ∼ DQB1*05:01, DQB1*03:03 ∼ DQB1*05:02, DQB1*05:01∼ DQB1*06:02,DQB1*05:01∼ DQB1*06:03, DQB1*05:01∼ DQB1*06:04, DQB1*05:02∼ DQB1*06:03, and DQB1*05:02∼ DQB1*06:04.

Considering the mean expression value of 365 in HLA-DPB1 locus, alleles DPB1*01:01 (n = 3), DPB1*03:01 (n = 14), and DPB1*14:01 (n = 3) were associated with a high expression, whereas alleles DPB1*02:01 (n = 11), DPB1*04:01 (n = 40), and DPB1*04:02 (n = 19) were associated with lower expression levels (Fig 8F). DPB1*05:01 (n = 4) was not linked to either due to its wide distribution of expression values. Different from the other loci, HLA-DPB1 showed a strinkingly heterogeneous distribution across the vast majority of alleles, excluding only DPB1*01:01, and hence no significant differences were found between different allele groups.

## Discussion

In the present study we demonstrate that it is possible to determine both the HLA alleles and their mRNA levels using RNA sequencing methodology. This type of tool can be applied in various approaches related to autoimmune and transplantation genetics as well as in studies of HLA expression levels in different cells and tissues, for example in the thymus. Despite the increasing evidence that HLA mRNA and surface protein expression differences may influence the immune response and susceptibility to several human diseases, only a few studies have systematically focused on the gene and especially the HLA allele-specific mRNA expression levels. The protein expression studies are certainly hampered by the fact that no allele-specific monoclonal antibodies recognizing all HLA alleles with equal affinity are available. Real-time PCR has been adopted in several studies for determining the expression of HLA alleles, however, the focus has mainly been on HLA class I.[3–5,10] Given the high number of known HLA alleles, real-time PCR approach requires a combination of allele-specific primers to amplify different alleles of the same locus. Using RNAseq data of 50 individuals, we performed a high-throughput screen for HLA expression profiles of class I and class II alleles in peripheral blood samples. To our knowledge, no method based on NGS has been reported for systematically quantifying the mRNA expression of HLA alleles.

Since genomic ONT data have been shown to be successful in HLA-typing [18,21], we explored the accuracy of ONT RNAseq data in HLA allele calling. The 2D reads from the full-length sequencing of HLA amplicons with MinION resulted in a good accordance with the Luminex reference methods at the 2-field resolution level, suggesting that HLA typing can be performed from targeted ONT RNAseq data. Our method provided a sufficient read depth for HLA class I and class II alleles to be assigned accurately with SeqNext-HLA. HLA class II genes showed more uniform distribution of read depth across the exons, whereas the coverage of HLA class I exon 1 and the beginning of HLA class I exon 2 were systematically lower in our data, independent from allele and gene. This may be due to a lower efficiency of reverse transcription enzyme with longer transcripts or a higher turnover of HLA class I mRNA. Moreover, this might have been the reason for the higher mismatch rate observed in HLA class I alleles since most of the polymorphisms lie in the exon 2 and 3 area. To ensure an adequate mRNA capture efficacy we chose the TSO’s UMI length to be 10 bp which we assumed still to provide sufficient complexity to enable corrections of PCR biases.

The comparison of allele ratios calculated from unique UMIs between the three datasets showed that both our targeted Illumina HLA amplicon and non-targeted Illumina cDNA method were able to quantitate the allele-specific expression differences. The same comparison between Illumina and ONT data, however, showed varying correlation values, suggesting that ONT is not yet able for accurate allele-level expression quantification. This is most likely due to the challenges of finding UMIs from the error-prone reads. A missing UMI position results in discarding the read leading to a reduced unique UMI count. Future improvements in the read quality could ease the UMI detection making ONT an option for HLA RNA sequencing. The comparison of Illumina datasets at the gene-level showed that HLA class II genes, and especially HLA-DR, were expressed at high levels in our targeted HLA amplicon data. This might be due to different efficacies of the gene-specific primers in the enrichment step or the fact that pooling of gene-specific PCR products was done in equal volumes instead of equal molarities. Even though our pipeline uses UMIs in PCR bias removal and considers only original transcripts, it is not able to correct bias between genes. Because Illumina cDNA method is not based on enrichment, we believe it is more accurate to quantify and compare the expression between genes as no bias is introduced in the library preparation step. Though, since the allele ratios were highly concordant between the two datasets, the targeted approach would be a valuable option for being more cost-effective. However, it still needs optimization in equalizing primer efficiencies and molarities between different HLA genes.

Although several HLA-typing tools for RNAseq data exist [23–25], they do not provide expression quantification with UMI counting. By using our custom pipeline we were able to determine HLA mRNA expression levels to the allele level. Our results of HLA class-level expression from cDNA data were concordant with previously reported [43] as HLA class I was expressed at higher levels than class II in all 50 samples. We also detected heterogeneity in the expression levels of HLA genes and heterodimers. Our results confirmed varying expression of HLA genes both within and between individuals. Despite a high interindividual variation, the data showed that HLA-B and HLA-C were equally abundant on transcript level and that they were expressed at higher levels than HLA-A. It is known that at the cell surface HLA-A and HLA-B are expressed at higher levels than HLA-C, however, is not entirely clear why this is. In a previous study low HLA-C protein level resulted from a faster degradation of HLA-C mRNA than HLA-A and HLA-B. [44] However, it is possible that HLA-C mRNA is initially levels similar to HLA-A and -B but post-transcriptional mechanisms such as inefficient assembly with β2-microglobulin affect its protein level expression. [44,45] Moreover, HLA-C mRNA expression can be tissue-dependent. In peripheral blood lymphocytes HLA-C had comparable mRNA levels to HLA-A and -B while in larynx mucosa it was lower.[46]

The imbalanced expression between HLA class II loci is in line with previous findings [43] as HLA-DR was confirmed to express at higher levels compared to HLA-DP and HLA-DQ. It is of note that we analysed the peripheral blood samples without any quantifications of their cellular contents and it is not clear how much variation in immune cell numbers affects the interindividual results.

To add one level of complexity we investigated the HLA allele-specific expression. Among our 50 samples we found distinct allele-specific expression profiles. This result has many interesting consequences worth further studies. For example, in the current transplantation donor selection only qualitative HLA allele typing is done. However, some previous studies have shown that the allele-level expression of a mismatched donor-recipient pair has an impact to the outcome of HSCT. [8,9] A mismatch between recipient’s high-expression allele and donor’s low-expression allele was found immunogenic and associated with an increased risk of acute GVHD and non-relapse mortality, whereas allotypes expressed at lower levels were not and hence were hypothesized as permissive. [8,9] In addition to the outcome of HSCT [8], differential expression of HLA class I genes or alleles have been associated with HIV control [6,12] and Crohn’s disease [11]. Considering the mean mRNA expression we were able to classify the alleles into high-expression and low-expression alleles. Among HLA-A alleles we found no significant difference between these two groups. However, our results showed that A*68:01 was expressed at higher levels compared to other HLA-A alleles and hence could be considered as immunogenic risk allele in HSCT and HIV control [12]. In contrast low-expression associated alleles such as A*01:01, and A*02:02, A*25:01, A*29:01, and A*29:02 with homogeneous expression distributions could be considered as possible permissive mismatches in HSCT. Our results are partly concordant with a previous study where the authors reported A*29 as an allele with a low expression.[4] However, in our data A*02:01 was associated with a lower mRNA expression demonstrating that the population origin can affect to the allele-specific expression. At HLA-B and HLA-C loci our results confirmed a significant difference in mRNA expression levels between high-expression and low-expression associated alleles indicating strong allele-specific expression. These loci showed more heterogeneous expression distributions within allele groups suggesting that the mRNA expression level is not always allele-bound. Due to the high haplotypic variety among our 50 samples, we did not inspect the effect of different haplotypes on HLA allele-specific expression. However, both HLA gene and allele level expression have shown to differ between haplotypes [3,47] and hence it is noteworthy that the heterogeneous expression within allele group might result from different haplotypes also in our data.

Variation in allele-specific expression of HLA-C has been already reported by a previous study.[3] Since our results are consistent with this data demonstrating C*01:02 and C*07:02 as low-expression associated alleles, and C*03:04 as high-expression allele, we can assume that some alleles are associated to high or low expression across populations, although this need further confirmation. HLA-C alleles, such as C*02:02, C*03:03, C*05:01, and C*06:02, were also linked to high expression levels. The risk allele of psoriasis [48], C*06:02, was observed to express at the highest level. These findings of the allele-specific expression are highly interesting from the perspective of human diseases. High HLA-C expression on cell-surface has already been shown to correlate with improved cytotoxic T lymphocyte response in HIV [6], as well increased risk for Crohn’s disease [11]. Moreover, the expression of HLA-C, which is the dominant ligand for natural killer (NK) cell killer immunoglobulin-like receptors (KIRs), was shown to associate with changes in NK subset distribution and licensing, especially in HLA-C1/C1, KIR2DL3+2DL2 individuals[49]. In addition to the enhanced T cell response, elevated HLA-C expression levels could affect NK cell development as well and result in a more effective respond upon infection.

The allele-level expression quantification also revealed differential expression profiles in class II genes. Despite heterogeneous expression profiles within allele groups, we observed HLA-DRB1 alleles associating with a high or low mRNA expression supporting the idea of allele-specific expression. The most striking differences in mean mRNA expression between alleles were seen at HLA-DRB3, and HLA-DRB5. In both genes the most frequent allele (DRB3*01:01, and DRB5*01:01) showed highest expression values and was dominated by hemizygous individuals. Since individuals carrying only one DRB3 or DRB5 allele were also expressed at lower levels, we concluded that there was no bias between hemizygous and heterozygous individuals in our data. However, we could not reliably determine allele-specific expression of HLA-DRA alleles. This locus turned out to be problematic for our pipeline as we observed a clear bias in unique UMI counts between heterozygous and homozygous individuals. We suspect that our pipeline could not quantify the allele-specific number of unique UMIs from Illumina short reads with the low number of polymorphic positions between HLA-DRA alleles. This is something we need to investigate further.

Our data showed a low allelic diversity at HLA-DPA1 with the majority of individuals carrying DPA1*01:03 which was a high-expression allele. DPA1*01:03 together with DPB1*04:02 has been reported as the most protective heterodimer from narcolepsy.[50] Considering the mean mRNA expression of HLA-DPB1 locus we found our results to be concordant with a previous study [9] associating alleles DPB1*01:01, DPB1*03:01, DPB1*14:01, and DPB1*15:02 to higher expression levels and alleles DPB1*04:01, and DPB1*04:02 to lower expression levels. However, it is notable that expression distributions at this locus varied greatly within several allele groups indicating that assigning alleles as high or low-expression linked is not straightforward. Interestingly, at HLA-DQB1 alleles DQB1*05:01 and DQB1*05:02 were expressed at clearly higher levels than the other HLA-DQB1 alleles. DRB1*01:01∼DQB1*05:01 haplotype was recently shown to be significantly protective for MS. [51] Moreover, DQB1*05:01 has been identified earlier as protective allele from narcolepsy [52,53] indicating that the high expression we see in our data would be beneficial at the population level. In contrast, the narcolepsy risk allele, DQB1*06:02 [54] and celiac disease risk alleles, DQA1*05:01, DQB1*02:01, DQA1*02:01, DQB1*02:02, HLA-DQA1*03, and DQB1*03:02 [55] were expressed at low levels.

Using RNAseq approach we have provided a new insight into the complexity of HLA allele-level expression. With increasing information of different factors affecting to the outcome of HSCT, it might be challenging to find a donor with suitable criteria and thus, make the donor selection more complicated. Therefore, our aim is to propose a tool to explore the differential HLA allele expression that in the future might ease the finding of possible permissive mismatches and help to avoid high-risk transplantations making HSCTs safer when no matched donor is available. Since several research and clinical HLA laboratories have already adopted NGS in HLA typing, the leap from DNA sequencing to RNAseq enabling both the HLA typing and expression quantification could be possible in the future changing the nature of HLA research from qualitative to quantitative.

## Materials and methods

### Samples and RNA extraction

This study collected 50 healthy blood donor buffy coat samples, which underwent an isolation of pheripheral blood mononuclear cells (PBMC) using Ficoll-Paque^™^ Plus (GE Healthcare), Dulbecco’s Phosphate Buffered Saline DPBS CTS^™^ (Gibco life technologies), Fetal Bovine Serum FBS (Sigma) and SepMate™-50 tubes following the manufacturer’s protocol (Stemcell Technologies). The use of anonymized PBMCs from blood donors is in accordance with the rules of the Finnish Supervisory Authority for Welfare and Health (Valvira). Cell count was measured from a mix of 50 µl of cell suspension in DPBS with 2% FBS, 50 µl of Reagent A100 lysis buffer, and 50 µl of Reagent B stabilizing buffer using a NucleoCassette and a NucleoCounter^®^ NC-100™ (all chemometec). Total RNA was isolated from fresh PBMC samples containing 1–10 x10^6^ cells using RNeasy Mini kit and Rnase-Free DNAse Set (both Qiagen) within two hours after PBMC isolation. RNA samples were quantified and the purity was assessed with the Qubit™ RNA High Sensitivity Assay Kit in Qubit^®^ 2.0 fluorometer (ThermoScientific). The RNA quality was checked using an RNA 6000 Pico Kit (Agilent Genomics) in a 2100 Bioanalyzer (Agilent Genomics) to obtain a RNA Integrity Number (RIN) score.

### Reverse transcription by template switching and target amplification

We used an adaptation of the STRT method to generate full length cDNA molecules from RNA transcripts.[31] Briefly, the poly-A hybridization to the first strand cDNA synthesis primer was performed in a 96-well plate in a T100™ Thermal Cycler (Biorad) with 3 min at 72°C with 25 ng of RNA, 1% Triton^™^ X-100 (Sigma), 20 µM of STRT-V3-T30-VN oligo, 100 µM of DTT (invitrogen, life technologies, Thermo Fisher), 10 mM dNTP (Bioline), 4 U of Recombinant RNase Inhibitor (Takara Clontech), 1:1000 The Ambion^®^ ERCC RNA Spike-In Control Mix (life technologies, Thermo Fisher) in a total volume of 3 µl. All oligos were from Integrated DNA Technologies and are listed in S1 Table. Reverse transcription of the whole transcriptome was performed adding 3.7 µl of the RT mix containing 5x SuperScript first strand buffer (invitrogen by Thermo Fisher Scientific), 1 M MgCl2 (Sigma), 5 M Betaine solution (Sigma), 134 U of SuperScript ^®^ II Reverse Transcriptase (invitrogen by Thermo Fisher Scientific), 40 µM RNA-TSO 10bp UMI, 5.6 U of Recombinant RNase Inhibitor immediately to each reaction. To complete the reverse transcription and the template switching the plate was incubated 90 min at 42°C followed by 10 min at 72°C. In this reaction every transcript receives a unique distinct barcode. After RT the cDNA was further amplified with 2x KAPA HiFi HotStart ReadyMix (Kapa Biosystems), 10 µM ImSTRT-TSO-PCR with a thermal profile consisted of an initial denaturation of 3 min at 95°C followed by 20 cycles of 20 s at 95°C, 15 s 55°C, 30 s at 72 and 1 cycle of final elongation of 1 min at 72°C in a final volume of 50 µl. Qubit™ dsDNA High Sensitivity Assay Kit (Thermo Fisher Scientific) was used to measure the concentration of all cDNA samples. The 3’ fragments of the cDNA were released in a restriction reaction using SalI-HH (New England Biolabs) according to the manufacturer’s protocol. The concentration of DNA was measured using Qubit™ dsDNA High Sensitivity Assay Kit and DNA integrity and the size distribution were assessed with High Sensitivity DNA Kit (Agilent Genomics). For HLA target enrichment one TSO-specific universal forward primer and eight gene-specific reverse primers with universal tails for amplicon sequencing were used to amplify exons 1 to 8 in class I genes HLA-A, -B, -C and -G or exons 1 to 5 in class II genes HLA-DRA, -DRB1, -DRB3, -DRB4, -DRB5, -DPA1, -DPB1, -DQA1 and -DQB1. HLA-A, -B and -C had one common primers as well as -DRB1, -DRB3, -DRB4 and -DRB5. All seven gene-specific primers were designed to fall within a non-polymorphic region using the known sequence diversity, as described in the international ImMunoGeneTics IMGT/HLA database (http://www.ebi.ac.uk/imgt/hla/). The amplification was performed in 96-well plates with 3 µl of template cDNA, 10x Advantage 2 PCR buffer, 50x Advantage^®^ 2 Polymerase Mix (Takara, Clontech), 10 mM dNTP (Bioline), 10 µM TSO forward primer and one of the seven HLA gene-specific reverse primers in a total volume of 15 µl. The PCR reaction consisted of an initial denaturation of 30 s at 98 °C following 3 cycles of 10 s at 98°C, 30 s at 55°C, 30 s at 72°C and 27 cycles of 10 s at 98°C, 30 s at 71°C, 30 s at 72°C and final elongation of 5 min at 72°C. To confirm the amplicon lengths and non-specific amplification 4 samples were selected from each plate with the amplification performed using different gene-specific primer. These samples were run on a 2% agarose gel (Bioline) with 10x BlueJuice^™^ loading dye (invitrogen by Thermo Fisher Scientific) in 0.5X TBE (Thermo Fisher Scientific) with the GelGreen™ (Biotium) and visualized using the Quick-Load 1kb DNA Ladder (New England Biolabs). DNA of the PCR amplicons was quantified with the Qubit™ dsDNA High Sensitivity Assay Kit and the fragment sizes analyzed with Agilent’s High Sensitivity DNA Kit.

HLA amplicons were pooled into two groups per sample by dividing genes that share the closest homology to different pools. The first pool contained genes HLA-A, -B, -C, -DRB1, -DRB3, -DRB4, -DRB5 and -DPB1 (henceforth gene pool 1) and the second HLA-DRA, -DPA1, -DQA1, -DQB1 and -G (henceforth gene pool 2). In the pooling 5 µl of PCR product was used from each PCR plate resulting in a final volume of 15 µl and 25 µl in gene pools 1 and 2, respectively. A purification and size selection of the pools were performed in a 0.7X beads:DNA ratio by using the Agencourt AMPure XP beads (Beckman coulter) according the manufacturer’s protocol and eluted in 15 µl of nuclease-free water. DNA of all 100 pools was quantified with the Qubit™ dsDNA High Sensitivity Assay Kit. The average fragment size distribution of gene pools 1 and 2 was assessed with Agilent’s High Sensitivity DNA Kit from 10 samples of both pools. The molarity of each pool was then calculated using the DNA concentration (ng/ µl) and the average fragment length (bp).

### ONT library preparation and sequencing

ONT sequencing compatible barcoded fragments were prepared in a PCR reaction 0.5 nM of DNA from gene pools, 2 µl of PCR barcode from the 96 PCR Barcoding Kit (ONT), 50 µl of LongAmp Taq 2x Mix (New England Biolabs) and Nuclease-Free water in a final volume of 100 µl where ONT’s universal tails were used as a template for barcode introducing primers. The PCR was performed in the following conditions; initial denaturation of 3 min at 95°C, following 15 cycles of 15 s at 95°C, 15 s at 62°C, 30 s at 65°C and a final extension step 3 min at 65°C. A second DNA purification and size selection was done in a 1X beads:DNA ratio by using the Agencourt AMPure XP beads according to the manufacturer’s instructions and eluted in 20 µl of nuclease-free water. After the purification DNA was quantified with the Qubit™ dsDNA High Sensitivity Assay Kit and barcoded PCR amplicons were pooled with equal molarities in 10 library pools in a total volume of 50 µl each consisting of 10 individuals and either 8 loci (gene pool 1) or 5 loci (gene pool 2). 1 µg of pooled barcoded PCR products were treated with the NEBNext Ultra II End-repair / dA-tailing Module (New England Biolabs) according a Ligation Sequencing Kit 2D (SQK-LSK208) protocol (ONT) using a DNA CS 3.6kb (ONT) as a positive control. A third DNA purification was performed using 1X beads:DNA ratio by using the Agencourt AMPure XP beads following the Ligation Sequencing Kit 2D protocol. ONT sequencing adapters were ligated using NEB Blunt / TA Ligase Master Mix (New England Biolabs) and Adapter Mix and HP Adaptor provided by ONT following a purification step using MyOne C1 Streptavidin beads (invitrogen by Thermo Fisher Scientific) according to the Ligation Sequencing Kit 2D protocol to capture HP adaptor containing molecules. The libraries were eluted in 25 µl of elution buffer and mixed with running buffer and library loading beads (ONT) prior to sequencing. All 10 libraries were sequenced for 48 hours on R9.4 SpotON flow cells (FLO-MIN106) on MinION Mk 1b device using the MinKNOW software (versions 1.1.21, 1.3.24, 1.3.25 and 1.1.30).

### Illumina library preparation and sequencing

For Illumina sequencing, all loci of 50 HLA amplicons were multiplexed per sample. 50 cDNA and 50 HLA amplicon libraries were prepared using the Nextera XT DNA Library Preparation Kit (Illumina). For an optimal insert size and a library concentration 600 pg of each cDNA and PCR amplicon sample was tagmented for 5 min at 55°C using 5 µl of Nextera’s Tagment DNA Buffer, 0.25 µl of Nextera’s Amplicon Tagment Mix in a final volume of 10 µl. The transposone was inactivated with 2.5 µl of Nextera’s Neutralize Tagment Buffer for 5 min at room temperature. The dual indexing and adapter ligation took place in a PCR reaction with 7.5 µl of Nextera PCR Master Mix, 4 µl of nuclease-free water and 10 µM of i5 custom oligo and 10 µM of Nextera i7 N7XX oligo using a limited-cycle PCR program: an initial denaturation 30 s at 95°C following 12 cycles of 10 s at 95°C, 30 s at 55°C, 30s at 72°C with a final elongation step of 5 min at 72°C. After the amplification all 50 cDNA and HLA amplicons samples were pooled together into two separate pools, one cDNA and one HLA amplicon pool. These two pools were then purified twice using the Agencourt AMPure XP beads according to the manufacturer’s instructions first with 0.6X beads:DNA ratio and then with 1X beads:DNA ratio and eluted in 30 µl. Qubit™ dsDNA High Sensitivity Assay Kit was used to quantify DNA and HT DNA HiSens Reagent kit and DNA Extended Range LabChip in LabChip GXII Touch HT (all PerkinElmer) to assess the size distribution of the libraries. A double size selection was performed with the Agencourt AMPure XP beads according to the manufacturer’s instructions to remove fragments over 1000 bp (0.8X beads:DNA ratio) and under 300 bp (0.6X beads:DNA ratio). Prior to sequencing the DNA concentration was assessed with Qubit™ dsDNA High Sensitivity Assay Kit HT DNA HiSens Reagent kit and the library size verified with HT DNA HiSens Reagent kit. The two pooled and barcoded libraries were denaturated with 0.2 M NaOH and diluted in the HT1 buffer to obtain a final library concentration of 20 pM in 0.95:0.05 cDNA:HLA amplicon ratio. The libraries were sequenced by using MiSeq and Nextseq sequencers with 600 cycles (Miseq v3) and 300 cycles (NextSeq 500/550 v2) kits (both Illumina) generating 300 bp and 150 bp pair-end sequence reads.

## Data analysis

ONT reads were processed using the 2D Basecalling plus barcoding for FLO-MIN106 250 bps workflow (version v1.125) on the cloud-based Metrichor platform (v2.45.5, v2.44.1, ONT) generating 1D template, 1D complement and 2D reads. The fastq files were extracted from the native fast5 files using NanoOK [32]. Illumina paired-end reads from cDNA and HLA amplicon libraries in fastq format underwent a UMI extraction using the UMI-tools (v0.5.11) [33] and were quality trimmed using trimmomatic (v0.35). HLA typing was done from ONT reads using SeqNext-HLA SeqPilot software (v.4.3.1, JSI Medical Systems) and Illumina Miseq reads using three different typing softwares: Omixon Explore (v1.2.0, Omixon), HLAProfiler [2], and an in-house HLA-typing tool (S1 Text). After this Miseq and Nextseq data were combined. Processed cDNA library reads were aligned using HISAT2 (v2.1.0) [34] to the human genome (GRCh38) and assigned to genes according to the UMI-tools pipeline using featureCounts tool from the subread package (v1.5.3) [35]. Samtools (v1.4) were used to sort and index BAM files and UMI-tools count tool to count the number of unique UMIs per gene. The set of 50 count files were then merged into a single count table using the Define NGS experiment tool in Chipster (v3.12.2) [36].

By using the allele types determined for each HLA gene, the reads of each sample were further processed to estimate their expression levels. The HLA genes are highly polymorphic, with more than 18,000 HLA alleles documented in the version 3.28.0 of IMGT/HLA reference database upon writing [37]. Despite the critical differences, the HLA gene sequences are highly similar resulting in very high multi-mapping of the reads. Thus, we implemented the strategy of assessing allele-specific expression by aligning reads, using last [38] only to selected reference sequences extracted from the IMGT/HLA HLA reference database.

For each HLA gene, all reads of a sample were aligned to a database containing only the reference sequences of the two identified alleles for the gene. For ONT reads, last was used with parameters -s 2 -T 0 -l 100 -a 100 -Q 1 for alignment of the template, complement and 2D reads. For Illumina reads, last with parameters -s 2 -T 0 -l 50 -a 100 -Q 1 -i1 was used for alignment of R1 reads only, R2 reads only, and paired end alignment (using last-pair-probs). The three Illumina read alignments were combined to include all reads that possibly originated from the two alleles. This alignment step filtered out reads that do not map to the two known alleles for the gene. The set of reads that aligned to the two references of the known alleles were retained, and their aligned portions along with their base qualities were extracted from the last MAF file format alignment output. To assign each read to either allele, (i) the polymorphic positions between the two reference sequences of the known alleles are identified by first performing multiple alignment of the two sequences (using msa R package) [39], and then getting the positions with high diversity (Shannon entropy index > 0.5) from the consensus matrix of the two sequences (generated using Biostrings v2.46.0 and ShortRead R packages) [40,41], (ii) the corresponding bases at the polymorphic positions are identified for the two reference sequences, (iii) reads from the set of retained reads that aligned only to either of the reference alleles, covering at least 30% of the polymorphic sites with at least 60% accuracy are kept (60% or more accurate matching at the polymorphic sites for the allele) and recorded as belonging to each allele; for reads from the set of retained reads that aligned to both alleles, their aligned portions are re-aligned separately to each reference allele sequence using overlap alignment (pairwiseAlignment function of Biostrings R package), then Bayesian statistical model is used to assign each read to either allele as follows: the read’s likelihood of originating from each of the two reference alleles is calculated based on how well the read matches the corresponding bases of the reference allele at the polymorphic positions, the likelihood is calculated as the sum of matches at the polymorphic positions given a reference allele (for a matching position, the match is quantified as the read base quality/maximum possible base quality, which is at maximum 1 for high quality bases in the read that match the reference allele base) divided by the number of polymorphic positions, a likelihood close to 1 suggests strong match between the read and the reference allele, the likelihoods of the read to the two reference alleles is calculated, the posterior probability for the two reference alleles given the read is then calculated by normalizing each likelihood by the sum of all likelihoods, the read is assigned to the reference allele with the higher posterior probability. Reads that cover less than 60% of the polymorphic sites between the two alleles are discarded. The remaining reads that are assigned to either allele are then combined with the previously recorded reads belonging to each allele from the previous step; for homozygous HLA genes, reads aligning to just one of the allele reference sequence that cover at least 30% of the polymorphic sites with at least 60% accuracy are kept, and (iv) to estimate allele-specific expression, all UMIs are extracted from the reads that belong to each allele. For Illumina reads, the UMIs are extracted from the read names. For ONT reads, the position of the TSO sequence is first pattern searched in the reads (using vcountPattern function of R Biostrings package), the 10 bases following the 3bp GGG at the end of the TSO sequence in the reads is extracted as the UMIs. Once all UMIs are collected for the reads belonging to an allele, UMIs are deduplicated by counting all UMIs within 1 Levenshtein distance (LD) only once. The total number UMIs after deduplication represent the expression of an allele.

After HLA expression quantification Illumina cDNA and HLA amplicon reads were normalized in three parts. First, HLA gene-specific counts resulting from the alignment of cDNA reads to the human genome were removed and replaced in the merged count table with HLA allele-specific UMI counts derived from cDNA reads after the custom pipeline. Second, read counts were normalized to counts per million (CPM) using the cpm tool from the limma package (v3.30.13)[42]. Third, number of unique UMIs of each allele in Illumina HLA amplicon libraries was normalized by calculating unique UMI proportions between alleles out of the total number of unique UMIs per sample. For each individual these proportions were then multiplied by the total number of CPM-normalized unique UMIs of all HLA alleles in cDNA library. To study the relationship between the class II transactivator (CIITA) and HLA class II expression, unique UMIs per CIITA were extracted from CPM-normalized cDNA data.

## Statistical Analyses

All statistical analyses were performed using non-parametric methods with GraphPad Prism v7.03 (GraphPad Software). The Spearman’s rank correlation and linear regression with 95% confidence intervals were applied in the comparison of allelic ratios between the datasets, and in the expression comparison of HLA class II and CIITA. Expression differences of heterodimer groups (HLA-A, -B, -C, -DR, -DQ, -DP) and HLA allele-specific expression (allele groups with n ≥ 3) were analyzed using the non-parametric Kruskal-Wallis test followed by the pairwise Dunn’s multiple comparisons test. For HLA class-level and gender-level comparisons pairwise analyses were performed using the Mann-Whitney U test. In all tests p-values < 0.05 were considered significant.

## Acknowledgements

We would like to thank Hanne Ahola and Sisko Lehmonen for their excellent technical assistance and personnel at Biomedicum Functional Genomics Unit (FuGU) and at national HLA laboratory at the Finnish Red Cross Blood Service. We would also like to thank CSC *–* IT Center for Science, Finland, for computational resources. This work was supported by grants from Clinical Research Funding (EVO/VTR), the Academy of Finland, and Tekes (the Finnish Funding Agency for Technology and Innovation).

---

**S1 Table. Primer sequences**

**S1 Text. HLA genotyping**

**S1 Fig. Experimental design of Illumina and ONT platform.**

In the library preparation process of Illumina and ONT mRNA is first transcribed into cDNA with simultaneous integration of 10 bp UMI in rnaTSO and further amplified. The full length cDNA is then divided and processed in parallel in Illumina’s and ONT’s protocol both involving an enrichment of HLA genes and adding sample-specific barcodes for multiplexing. In Illumina’s protocol both full length cDNA and HLA amplicons are tagmented resulting in 5’ end library molecules.

**S2 Fig. Comparison of the number of raw reads between Illumina and ONT MinION datasets according to 50 individuals.**

White bars correspond to Illumina cDNA reads, grey bars to Illumina HLA amplicon reads, and black bars to barcoded ONT reads. The ONT sequencing of gene pools 1 and 2 on SpotON flow cells with the R9.4 chemistry generated 22,487 to 193,467 barcoded reads per sample. Illumina sequencing of the tagmented cDNA and HLA amplicons on MiSeq and Nextseq in total generated 497,134 to 6,649,598, and 36,638 to 169,116 reads per sample, respectively.

**S3 Fig. HLA typing accuracy of ONT dataset and concordance with Luminex.**

(A–B) The concordance rates of SeqNext-HLA typing results from ONT and Illumina datasets and at 1-field and 2-field resolution level. Alleles assigned by SeqNext-HLA were 100% concordant at 1-field level with alleles assigned by Luminex. At 2-field level the allele assigned by SeqNext-HLA was considered concordant if it was found in the list of alleles by Luminex technology. HLA-DRB1, -DRB3, -DRB5 and -DPB1 were 100% concordant with Luminex and with HLA-A, -B, -C, -DRB4, -DQA1, -DQB1 and -DPA1 the concordance rate was between 94% and 99%. No reads were assigned to the HLA-G gene. (C) Gene-specific distribution of mismatches between the allele assigned by SeqNext-HLA and the closest reference allele. (D–E) The concordance rates of ensemble typing results and Luminex HLA typing at 1-field and 2-field resolution level. At 1-field level all loci but HLA-DQB1 were over 90% concordant with the reference alleles. At 2-field the concordance rate for HLA-A, -B, and -C was 95%, 87%, and 86%. In class II the concordance rate varied from 71 to 99%. With Illumina data, in case of an expression difference within a heterozygous allele pair, the second allele was sometimes missed and the genotype was falsely assigned as homozygous.

**S4 Fig. The proportion of total and class-level HLA expression of the whole transcriptome expression according to 50 individuals.**

(A) Total HLA expression was calculated from normalized unique UMI counts of all HLA genes per individual and dividing this sum by the total number of normalized unique UMIs of the whole transcriptome. The percentages of HLA class I (B) and HLA class II (C) were calculated in a similar manner.

**S5 Fig. The comparison of HLA class I allele-specific expression values between Illumina amplicon and Illumina cDNA data.**

The expression profiles showing the normalized allele-level unique UMI counts of HLA class I genes (A– B) HLA-A, (C–D) HLA-B, (E–F) HLA-C in Illumina amplicon and cDNA data according to the 50 individuals. Mean expression of individual alleles is indicated by a solid bar and mean expression of all alleles is represented by the dotted line. Open circles correspond to homozygous individuals.

**S6 Fig. The comparison of HLA class II allele-specific expression values between Illumina amplicon and Illumina cDNA data.**

The expression profiles showing the normalized allele-level unique UMI counts of HLA class II genes (A–B) HLA-DRB1, (C–D) HLA-DRB3, HLA-DRB4, HLA-DRB5, (E–F) HLA-DPA1, (G–H) HLA-DPB1, (I–J) HLA-DQA1, (K–L) HLA-DQB1 of 50 individuals according to alleles. Mean expression of individual alleles is indicated by a solid bar and mean expression of all alleles is represented by the dotted line. Open circles correspond to homozygous individuals and black triangles to hemizygous individuals.

**S2 Table. UMIs from Illumina cDNA data.**

**S3 Table. UMIs from Illumina amplicon data.**

**S4 Table. UMIs from Nanopore data.**

